# A glycoprotein mutation that emerged during the 2013-2016 Ebola virus epidemic alters proteolysis and accelerates membrane fusion

**DOI:** 10.1101/2020.07.13.201863

**Authors:** J. Maximilian Fels, Robert H. Bortz, Tanwee Alkutkar, Eva Mittler, Rohit K. Jangra, Jennifer S. Spence, Kartik Chandran

## Abstract

Genomic surveillance of viral isolates during the 2013-2016 Ebola virus epidemic in Western Africa—the largest and most devastating filovirus outbreak on record—revealed several novel mutations. The responsible strain, named Makona, carries an A to V substitution at position 82 in the glycoprotein (GP), which is associated with enhanced infectivity *in vitro*. Here, we investigated the mechanistic basis for this enhancement, as well as the interplay between A82V and a T to I substitution at residue 544 of GP, which also modulates infectivity in cell culture. We found that both 82V and 544I destabilize GP with the residue at 544 impacting overall stability, while 82V specifically destabilizes proteolytically cleaved GP. Both residues also promote faster kinetics of lipid mixing of the viral and host membranes in live cells, individually and in tandem, which correlates with faster times to fusion following co-localization with the viral receptor Niemann-Pick C1 (NPC1). Further, GPs bearing 82V are more sensitive to proteolysis by cathepsin L (CatL), a key host factor for viral entry. Intriguingly, CatL processed 82V variant GPs to a novel product of ∼12K size, which we hypothesize corresponds to a form of GP more fully primed for fusion than previously detected. We thus propose a model in which 82V promotes more efficient GP processing by CatL, leading to faster viral fusion kinetics and higher infectivity.

**Importance:** The 2013-2016 outbreak of Ebola virus disease in West Africa demonstrated the potential for previously localized outbreaks to turn into regional, or even global, health emergencies. With over 28,000 cases and 11,000 confirmed deaths, this outbreak was over 50 times as large as any previously recorded. This outbreak also afforded the largest ever collection of Ebola virus genomic sequence data, allowing new insights into viral transmission and evolution. Viral mutants arising during the outbreak have attracted attention for their potentially altered patterns of infectivity in cell culture, with potential, if unclear, implications for increased viral spread and/or virulence. Here, we report on the properties of one such mutation in the viral glycoprotein, A82V, and its interplay with a previously described polymorphism at position 544. We show that mutations at both residues promote infection and fusion activation in cells, but that A82V additionally leads to increased infectivity under cathepsin-limited conditions, and the generation of a novel glycoprotein cleavage product.

## Introduction

Ebola virus (EBOV), a member of the viral family *Filoviridae*, was responsible for the largest recorded outbreak of filovirus disease in history, in Western Africa from 2013–2016. This epidemic is associated with over 28,000 suspected or confirmed cases and caused over 11,000 deaths. Genomic surveillance of viral isolates from infected patients have yielded a wealth of information on how the virus spread and evolved during the epidemic. The index case is thought to be a four-year old boy in Guinea, who may have been exposed to EBOV as a result of contact with an infected bat. The virus then spread to neighboring Liberia and Sierra Leone in 2014 exclusively through human-to-human transmission (1–4). The EBOV strain responsible for this outbreak was named Makona (5) and is genetically distinct from the Mayinga isolate from the first recorded EBOV outbreak (Democratic Republic of Congo, 1976).

Three missense mutations in the EBOV Makona genome sequence, (i) R111C in the nucleoprotein (NP), (ii) D759G in the polymerase (L), and (iii) A82V in the viral glycoprotein (GP), arose early in the epidemic and came to dominate the viral isolates collected during the last half of 2014 and onward (6). The A82V mutation in particular was proposed to be an adaptation arising during the prolonged chain of human-to-human transmission. This is supported by an observed increase in infection of primate cells and a corollary decrease in infection of bat cell lines for viruses carrying GP(A82V) (7, 8). Although this observation has not been recapitulated in animal models of infection, including mice and rhesus macaques *(9)*, there is mechanistic evidence that the A82V mutation impacts viral entry (10). Entry of EBOV depends on several host factors, including the obligate intracellular filovirus receptor Niemann-Pick C1 (NPC1) (11, 12), as well as cysteine cathepsin proteases that cleave GP to reveal the receptor-binding site (RBS) (13, 14). It remains unknown if the A82V mutation promotes more efficient usage of any of these known host factors, or if it allows the virus to co-opt novel host factors in order to mediate entry.

Another polymorphism in EBOV GP, at position 544, has also been shown to influence viral entry in tissue culture, with 544I conferring enhanced infectivity relative to 544T (15–18). Because Makona isolates have 544T while Mayinga isolates have 544I, we postulated that the A82V mutation in Makona arose, at least in part, as an infection-promoting response to the presence of 544T.

Here, we set out to investigate the interplay between polymorphisms at positions 82 and 544 in the background of GP(Makona) and GP(Mayinga). We found that both 82V and 544I mutations accelerated the kinetics of viral membrane fusion in an additive manner. Further, 82V variants were more susceptible to proteolysis by the endosomal cysteine protease cathepsin L (CatL) and displayed a striking alteration in the pattern of GP proteolytic cleavage, which correlated with enhanced viral entry under certain conditions. Taken together, our findings reveal a potential mechanism by which the A82V mutation enhances EBOV GP-dependent viral entry and infection.

## Results

### GP variants recapitulate previously reported infection and receptor binding phenotypes

We generated four GP variants - Makona 82A/544T, Makona 82V/544T, Mayinga 82A/544I, and Mayinga 82V/544I [henceforth referred to as GP(Mak 82A/544T), GP(Mak 82V/544T), GP(May 82A/544I), and GP(May 82V/544I), respectively] as vesicular stomatitis virus (VSV) pseudotypes (VSV-GP). Since none of the mutations of interest are in the GP mucin domain, and the mucin domain is dispensable for infection in cell culture (14, 19), all pseudotypes generated have a deletion of amino acid residues 309–489 in GP.

In order to validate these pseudotyped viruses, we first measured their infectivity in Vero cells. The A82V and T544I mutations, both alone and together, promoted higher levels of infectivity (**Figure 1A**), as previously reported (15–18). The differences in infectivity did not stem from differences in GP density on the viral surface, as all four variants demonstrated comparable levels of GP incorporation into particles (**Figure 1B**).

**Figure 1.**
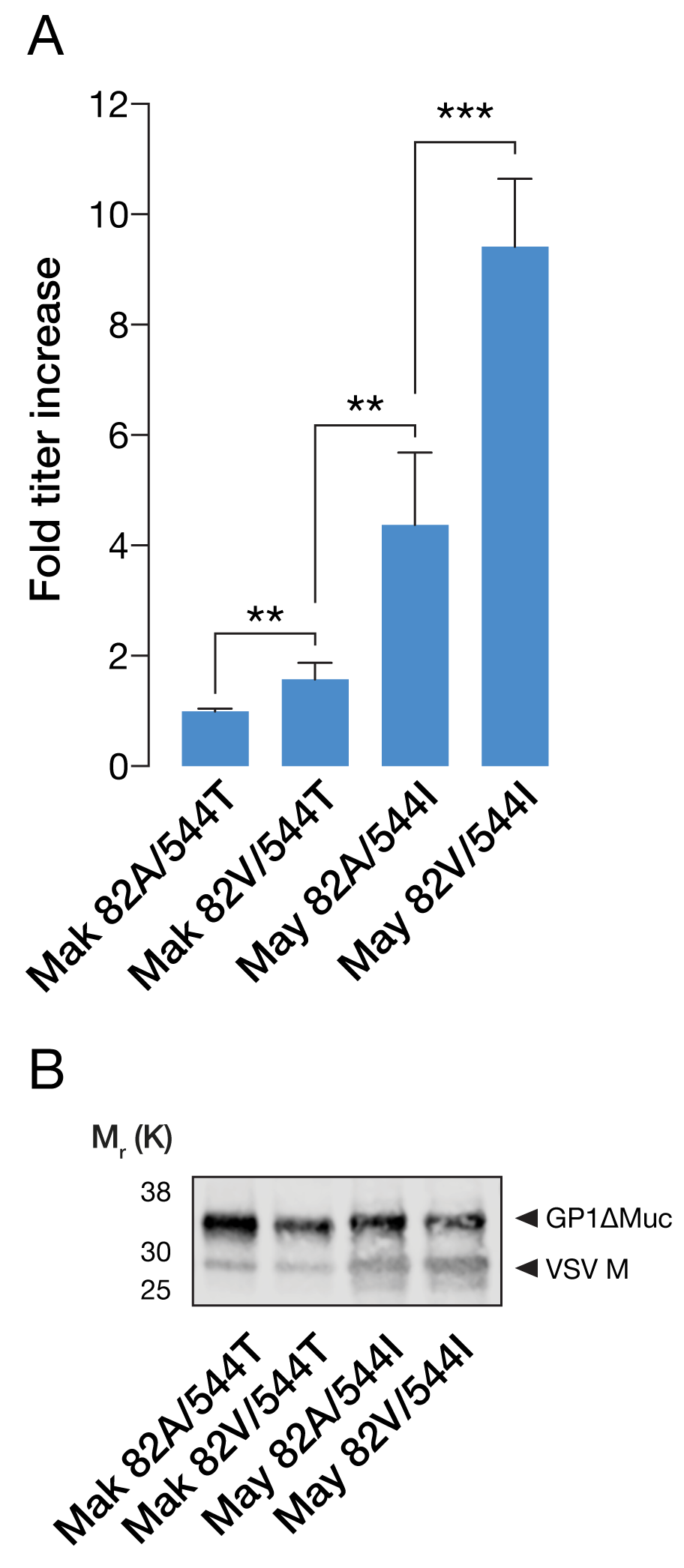
**(A)** Mean fold-increase in viral titers (+/- SD) of GP variants as compared to GP(Mak 82A/544T). 82V, 544I, or both in tandem, significantly increase titers (**, p < 0.002, ***, p < 0.001, by unpaired two-tailed *t*-tests) (n = 6, from three independent experiments). **(B)** Representative western blot of VSV M and EBOV GP indicating similar levels of GP incorporation into VSV for all four GP variants.

The various infectivity phenotypes of the mutants also could not be readily explained by differences in receptor interaction. Due to the proximity of position 82 to the RBS of GP, it has been speculated that the A82V mutation could promote binding to the critical endo/lysosomal filovirus receptor, Niemann-Pick C1 (NPC1) (11, 12, 20) (7). Following pre-treatment of the VSV-GPs with thermolysin (THL), which mimics the activity of cathepsin B (CatB) (14), we used an enzyme-linked immunosorbent assay (ELISA) to measure NPC1 domain C binding to cleaved GP (GP_CL_) (21). As reported previously (10), the A82V mutation did not affect GP_CL_-NPC1 binding (**Figure 2**).

**Figure 2.**
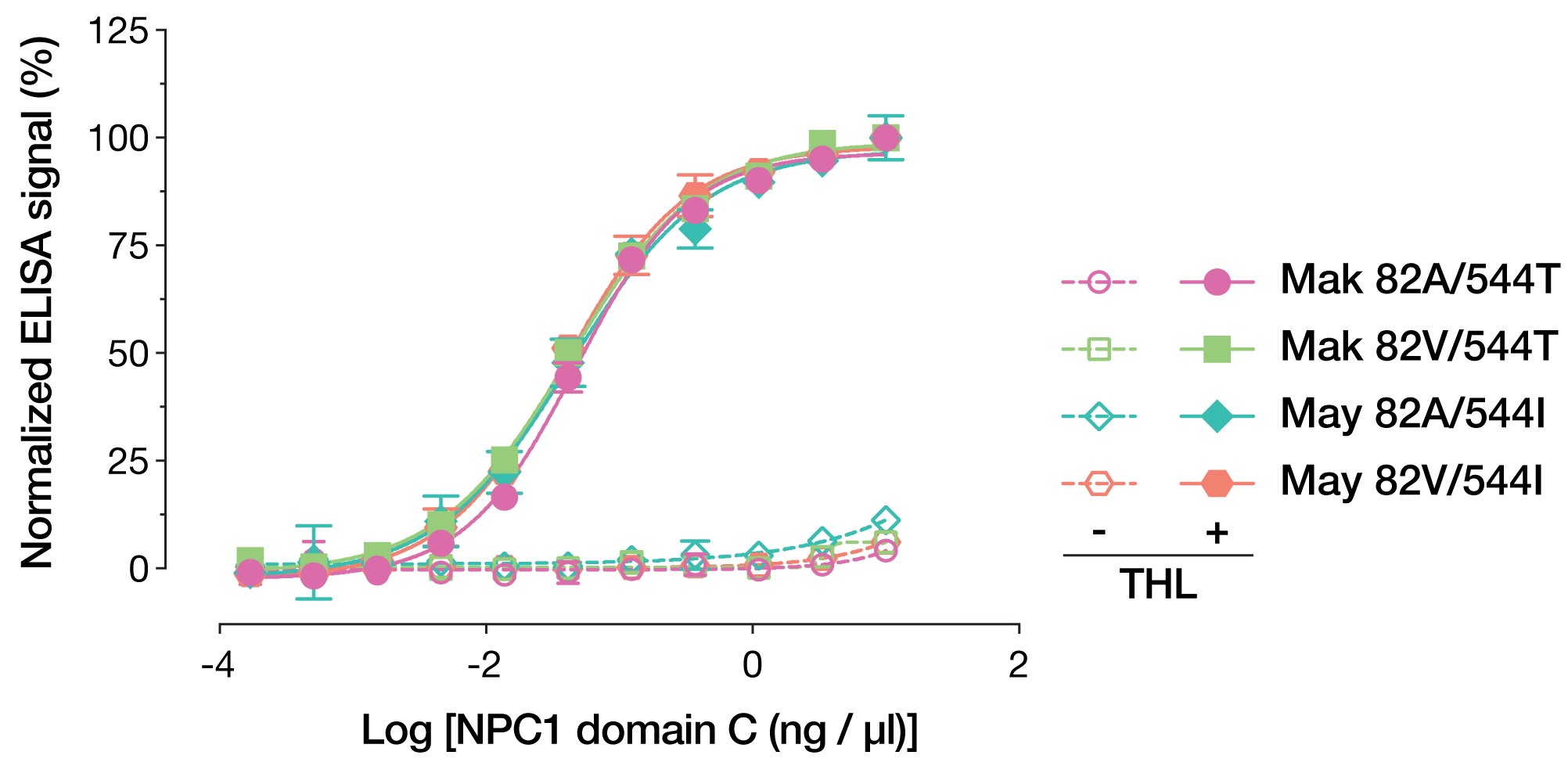
Normalized amounts of VSV-GP variants, in native or THL-cleaved state, were captured onto ELISA plates using KZ52. Once cleaved, all four GP variants are able to bind NPC1 domain C at equal levels. Means (+/-SD) of results from two independent experiments with three technical replicates are shown (n = 6).

One of the most striking and widely reported phenotypes associated with the A82V mutation is increased resistance to the small-molecule viral inhibitor 3.47, which targets the GP-NPC1 interaction (12, 22). We observed that the polymorphisms at positions 82 and 544 made independent and additive contributions to viral sensitivity to 3.47. Specifically, A82V and T544I were each associated with increased 3.47 resistance, with the 82A/544T and 82V/544I genotypes conferring low and high resistance, respectively, and the mixed 82V/544T and 82A/544I conferring intermediate levels of resistance (**Figure 3**). These results are largely in line with previously reported data (10) and indicate that both A82V and T544I promote viral resistance to 3.47 in a manner likely unrelated to GP-NPC1 interaction.

**Figure 3.**
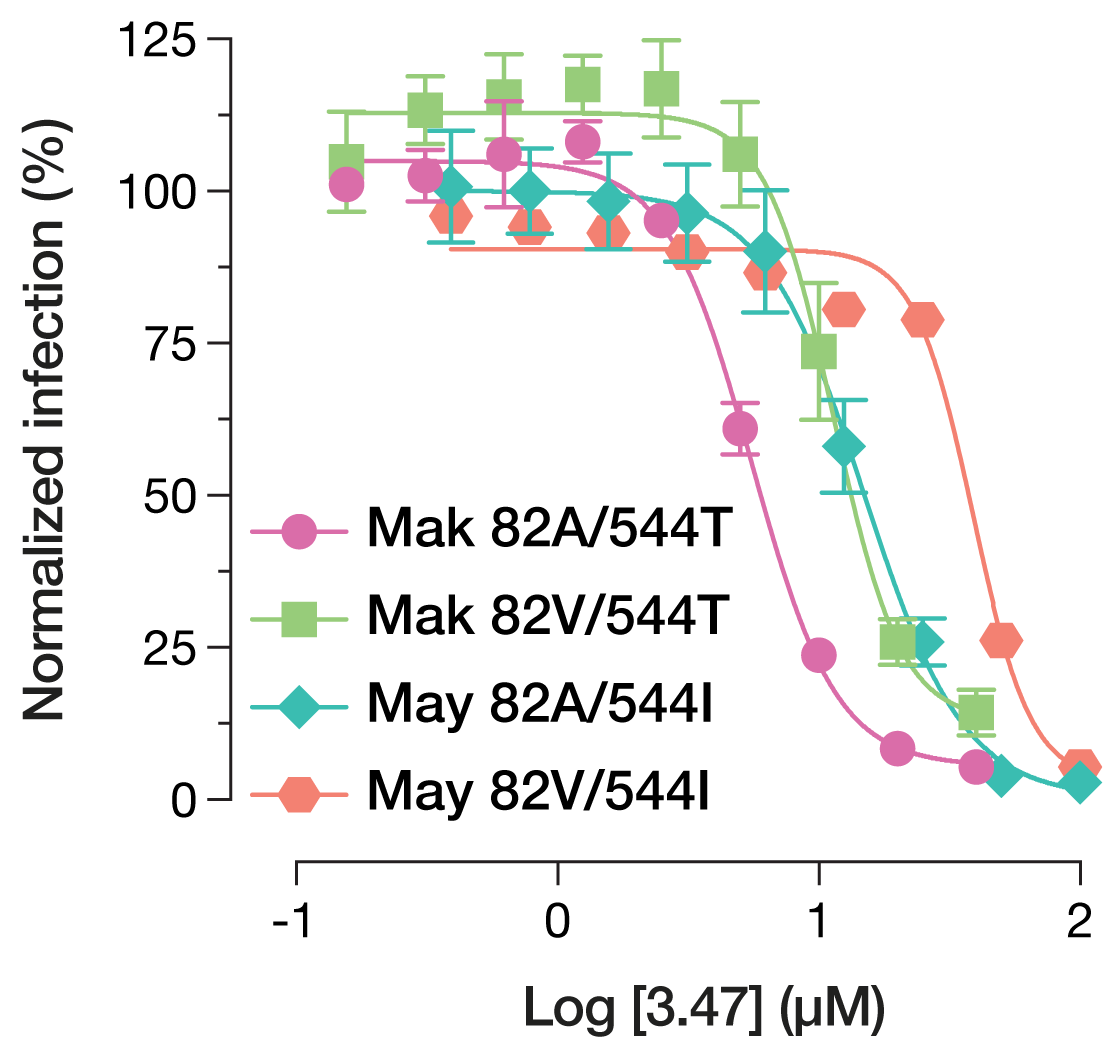
Normalized infectivity levels at varying concentrations of the viral inhibitor 3.47. Means (+/-SD) of results from two independent experiments with three technical replicates are shown (n = 6). At 3.47 concentrations >2.5 and >12.5 µM, respectively, Makona and Mayinga VSV-GP variants bearing 82V are significantly more infectious than their 82A counterparts (p < 0.001, by multiple t tests with the Holm-Šídák correction for multiple testing).

### The GP destabilizing effect of the A82V mutation is unmasked by proteolysis

Single point-mutations have previously been shown to have dramatic effects on GP stability with associated impacts on viral infectivity (23). A previous report implicated the reduced stability of the A82V mutant as the basis for its increased infectivity *in vitro* and *in vivo* (10). Likewise, proteolysis has a destabilizing effect on GP that is thought to promote fusion activation (23–25). Accordingly, we set out to investigate the stability of GP mutants in both uncleaved and pre-cleaved states, using an ELISA that measures the thermostability of an epitope spanning GP1 and GP2 (25). By measuring the half-maximal melting temperature (T_m_) of GPs, we found that uncleaved GPs bearing 544I were significantly less stable than their 544T counterparts (**Figure 4A and 4C**). Upon THL cleavage, all GPs were destabilized, with T_m_ values decreasing 5–6°C compared to their uncleaved counterparts. Further, THL cleavage unmasked more nuanced differences in stability, with GPs bearing 82V being significantly destabilized as compared to the 82A variants (**Figure 4B and 4D**). Thus, the polymorphisms at positions 82 and 544 both appear to regulate GP stability, but may do so at different steps in entry—either upstream (544) or downstream (82) of the proteolytic cleavage mimicked by THL.

**Figure 4.**
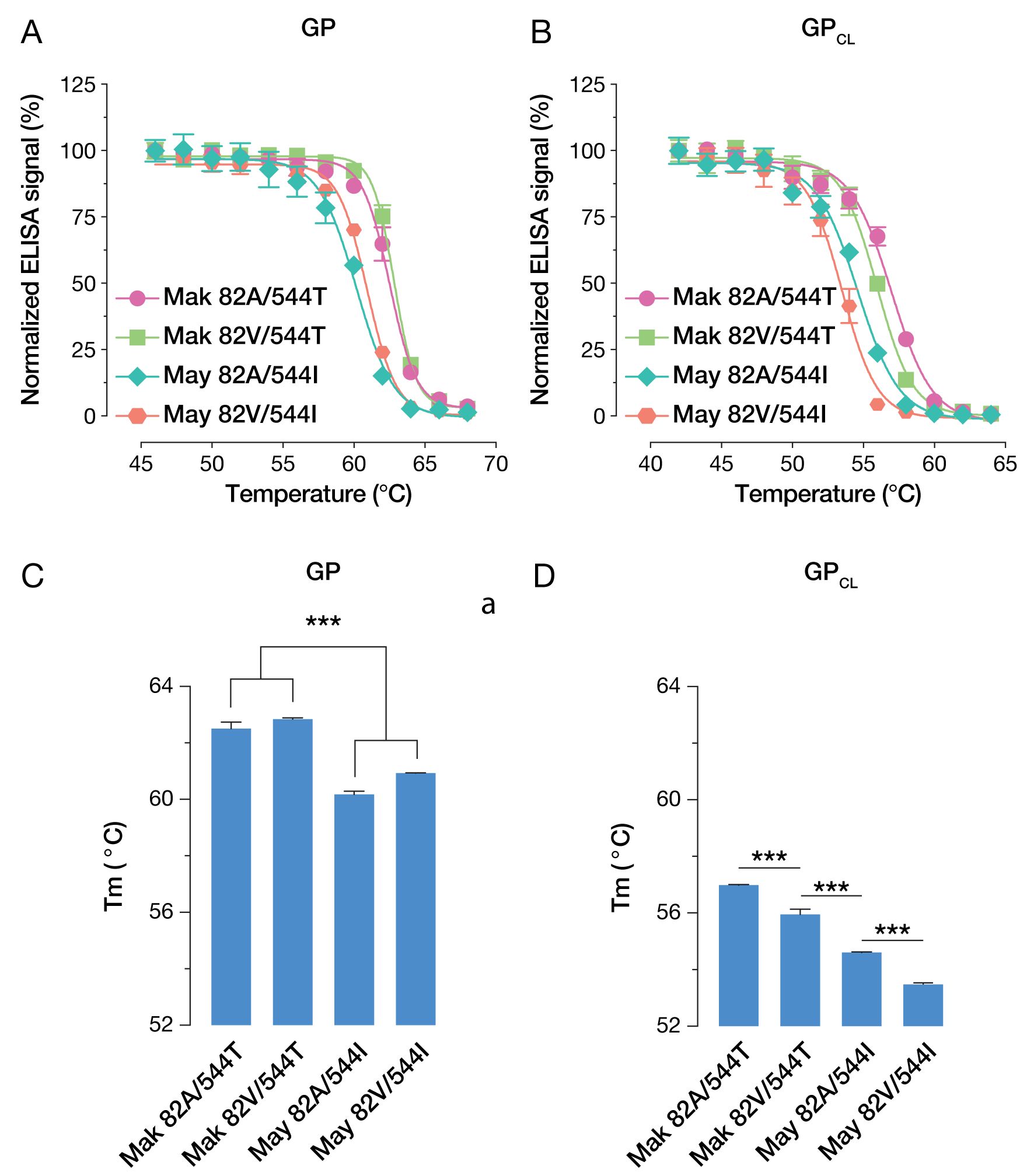
**(A)** Normalized ELISA signal of KZ52 binding to native VSV-GP variants following heating. Means (+/-SD) of results from two independent experiments with three technical replicates (n = 6) are shown. **(B)** Normalized ELISA signal of KZ52 binding to THL-cleaved VSV-GP variants following heating. Means (+/-SD) of results from two independent experiments with three technical replicates (n = 6) are shown. **(C)** T_m_-values for native VSV-GP variants, derived through non-linear regression analysis of ELISA signals. Statistical significance was determined by one-way ANOVA with Tukey correction for multiple testing (*, p <0.033; **, p <0.002; ***, p <0.001) **(D)** T_m_-values for THL-cleaved VSV-GP variants, derived through non-linear regression analysis of ELISA signals. Statistical significance was determined by one-way ANOVA with Tukey correction for multiple testing (*, p <0.033; **, p <0.002; ***, p <0.001).

### A82V and T544I increase the rate of GP fusion activation but not its probability

In order to investigate the impact of these GP mutations on fusogenic activation, we used a fluorescence dequenching assay to observe viral fusion triggering within live cells on a single-particle basis (26, 27). Membranes of purified viruses were labeled with self-quenching concentrations of the lipophilic dye DiD. Mixing of lipids between the host and viral membranes as a result of hemifusion or full fusion enables dequenching of the dye and a sharp increase in fluorescence intensity. We observed that the polymorphisms at positions 82 and 544 made independent and additive contributions to the kinetics of fluorescence dequenching (**Figure 5A**). Specifically, VSV-GP(May 82V/544I) exhibited the fastest kinetics of lipid mixing, with a *t*_1/2_ (time needed for 50% of particles to dequench) of 24 min, whereas VSV-GP(Mak 82A/544T) exhibited the slowest (*t*_1/2_ = 57 min). Viruses with the mixed GP genotypes exhibited intermediate kinetics of lipid mixing (*t*_1/2_ = 44 and 47 min for 82A/544I and 82V/544T, respectively). Interestingly, the lipid mixing kinetics of these viruses in cells were concordant with their respective susceptibilities to inhibition by 3.47 (**Figure 3**).

**Figure 5.**
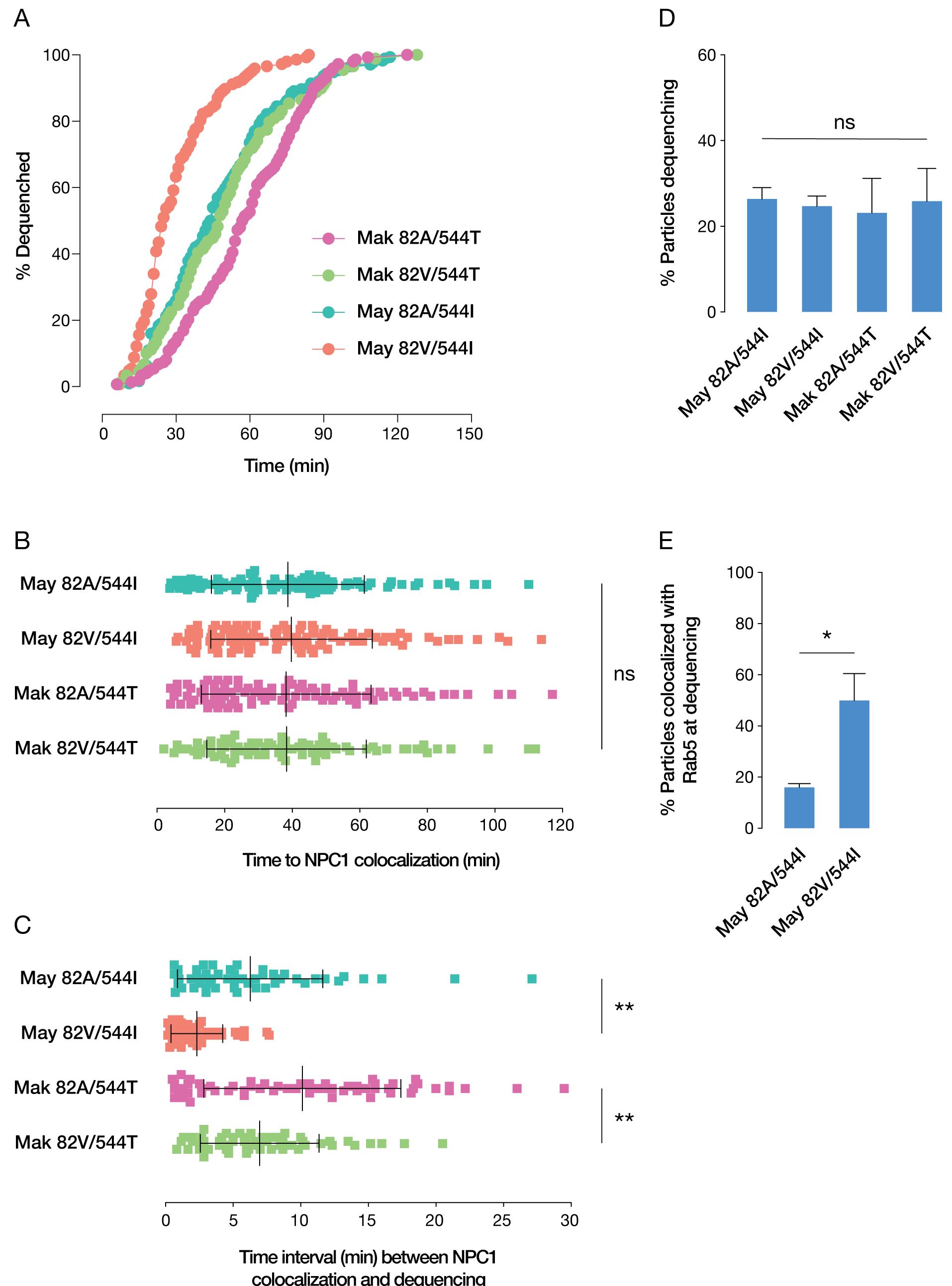
**(A)** Kinetics of viral fusion triggering. **(B)** Time of viral colocalization with NPC1. Data represent the mean and SD of three independent experiments. **(C)** Time interval between viral colocalization with NPC1 and onset of lipid mixing. Data represent the mean and SD of three independent experiments (** p < 0.01 by one-way ANOVA with a post-hoc Tukey test). **(D)** Total percentage of virions undergoing GP-mediated lipid mixing. Data represent the mean and SD of three independent experiments. **(E)** Total percentage of virions colocalized with Rab5 at the time of lipid mixing (* p < 0.05 by Student’s *t* test).

Despite these differences in the kinetics of GP-dependent lipid mixing in cells, however, we observed no difference with respect to the total percentages of cell-associated particles eventually inducing lipid mixing (**Figure 5D**). Since this assay is unable to differentiate hemifusion from the formation of fusion pores and viral content release into the cytoplasm, this suggests that the differences in infectivity observed for the 82 and 544 mutants arise at a very late step in fusion activation that we were unable to resolve other than as a change in the kinetics of fusion activation. It is also unknown how the timing of fusion impacts the probability to establish productive infection, adding a layer of complexity to inferences about the relationship between lipid mixing and infectivity.

We also observed no significant differences among the mutants in terms of mean time of colocalization with NPC1, demonstrating that internalization and trafficking are unaltered (**Figure 5B**). However, once delivered to NPC1-positive compartments, VSV-GP(May 82V/544I) exhibited a shorter mean lag to lipid mixing (2.3 min) versus GP(May 82A/544I) (6.3 min) (**Figure 5C**). The lag increased slightly with GP(Mak 82V/544T) (6.9 min) and to a greater extent with GP(Mak 82A/544T) (10.1 min).

Because the dramatically accelerated fusion kinetics of GP(May 82V/544I) indicated that its threshold for triggering may be significantly reduced, we examined lipid mixing in Rab5-positive compartments (**Figure 5E**). The percentage of virions undergoing dequenching in intermediate endosomes, as differentiated from early endosomes by the presence of NPC1 at the time of fusion, was significantly higher with this mutant versus GP(May 82A/544I), demonstrating that more particles can be triggered to fuse in less-mature endocytic vesicles.

### 82V variants have greater relative infectivity in CatL-deficient cells

To account for the differing fusion kinetics observed among the GP variants, we next examined GP proteolytic susceptibility *in vitro*. Previous findings indicate that both the A82V and T544I mutations influence the CatB dependence of viral entry. While T544I alone does not promote resistance to CA074, the residue at position 544 does influence the overall dependence on CatB (10, 18). Here, we found that only the 82V/544I variant was resistant to the CatB inhibitor CA074 (**Figure 6A**), largely consistent with Wang and co-workers’ observations. Because CatL can support EBOV GP-dependent entry (13, 14, 28), we next sought to investigate the effect of the 82/544 polymorphisms on viral entry under CatL-limited conditions. To overcome the challenges associated with selective inhibition of CatL in cells with irreversible activity-based inhibitors, we previously generated U2OS CatL KO cells through CRISPR/Cas9 genome engineering (29). We then exposed control and CatL KO cells to VSVs bearing the Makona and Mayinga 82/544 GP variants. Both 82V variants afforded higher infectivity than their 82A counterparts in CatL-KO cells even after adjusting for their relative infectivities in the control cells (**Figure 6B**). Taken together, our results suggest that although A82V and T544I are required to bypass the entry requirement for CatB, A82V but not T544I enhances viral entry in the absence of CatL.

**Figure 6.**
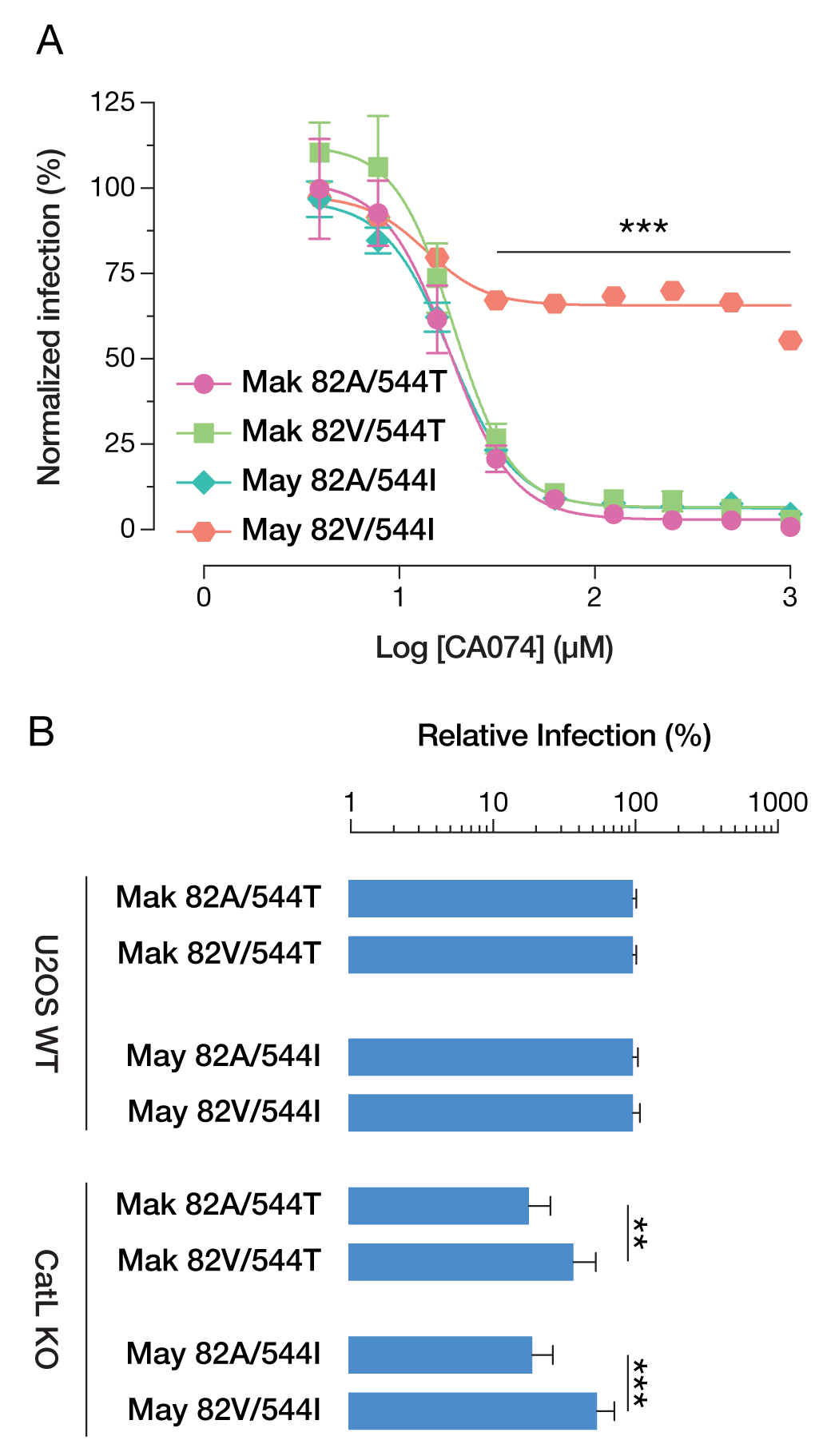
**(A)** Normalized infectivity levels at varying concentrations of the CatB inhibitor CA074. Means (+/-SD) of results from two independent experiments with three technical replicates (n = 6) are shown. At concentrations of CA074 >15 µM, VSV-GP(May 82V/544I) is significantly more infectious than all other GP variants (P < 0.001, by multiple *t* tests with the Holm-Šídák correction for multiple testing). **(B)** WT and CatL KO U2OS cells were infected with VSV-GP variants, and infectivity was scored by automatic counting of GFP-positive cells 14 hours post-infection. Groups were compared by one-way ANOVA with Šídák correction for multiple testing in order to determine statistical significance (*, p <0.033; **, p <0.002; ***, p <0.001). On CatL KO cells, viruses carrying 82V variants have a higher relative infectivity than those carrying 82A. Means (+/-SD) of results from nine trials from three independent experiments with three technical replicates are shown.

### A82V mutations allow GP to be processed to a novel cleavage product by CatL

To explore the molecular basis of enhanced viral entry in CatL-deficient cells afforded by the A82V mutation, we subjected VSVs bearing all four GP(82/544) variants to *in vitro* proteolytic cleavage with CatL or THL (to mimic CatB activity) and analyzed the deglycosylated cleavage products by SDS-PAGE (**Figure 7A**). With CatL treatment, we observed a rapid conversion to an 18K species (GP1_18K_) for all four GP variants. GPs with 82A were resistant to further proteolysis, as evidenced by the persistence of GP1_18K_ for up to 3 h. GPs with 82V, by contrast, were converted to a novel GP1 cleavage product of approximately 12K size (GP1_12K_). Both GP1_18K_ and GP1_12K_ were degraded upon prolonged CatL treatment. Treatment with THL led only to the formation of the expected 17K product (GP1_17K_) for all four GP variants. No overt differences in susceptibility to THL were observed in 1 h of THL treatment. Therefore, the presence of 82V appears to induce a specific change in GP sensitivity to CatL rather than a generalized change in its sensitivity to proteolysis (**Figure 7B**).

**Figure 7.**
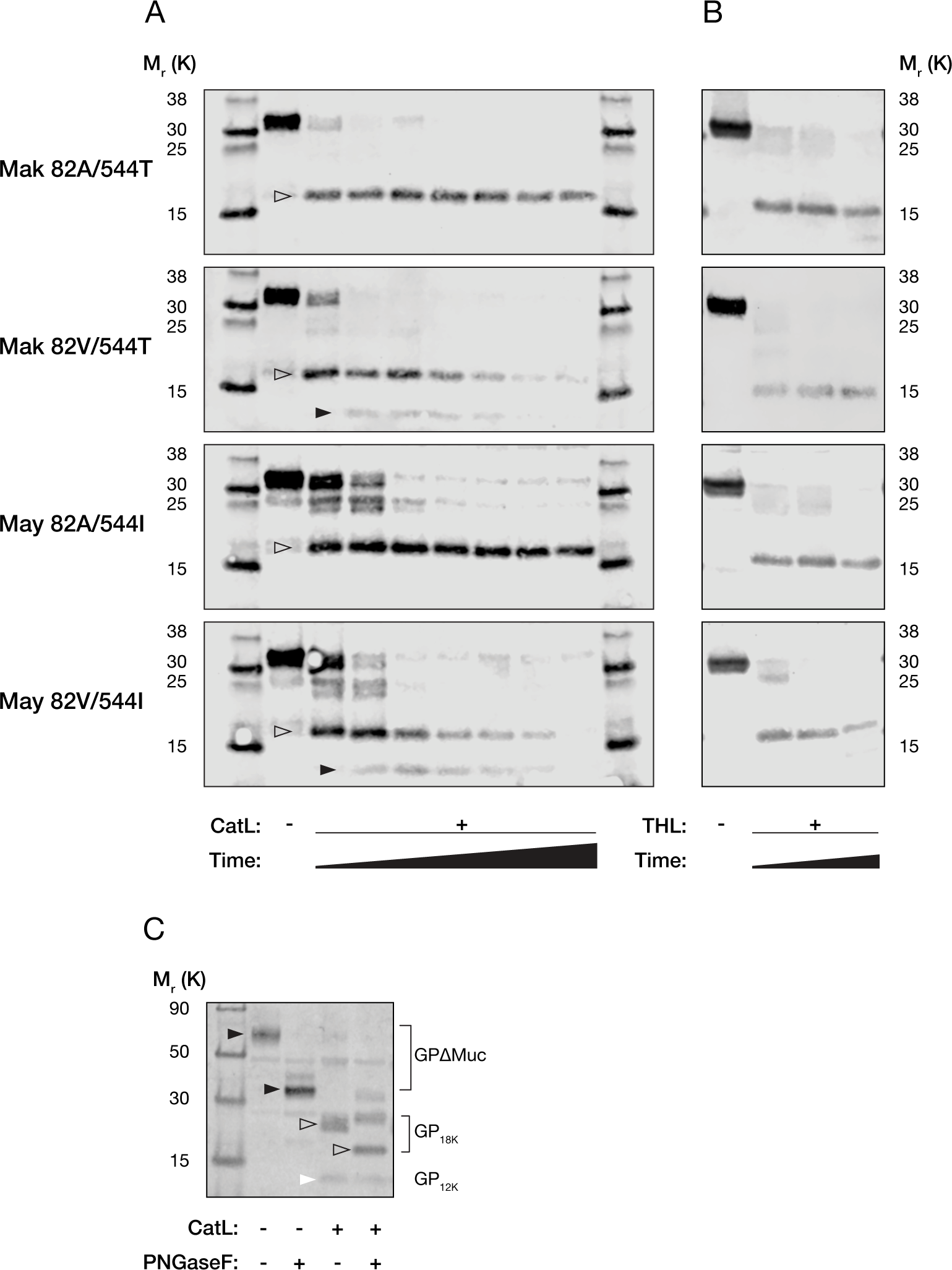
**(A)** Representative western blots of GP1 CatL cleavage products analyzed by SDS-PAGE following PNGase F treatment. With increased incubation times (0 - 3 h) GP variants bearing 82V were converted to two distinct products of approximately 18K and 12K, respectively. **(B)** Representative western blots of GP1 THL cleavage products analyzed by SDS-PAGE following PNGase F treatment. No overt differences in proteolysis were observed between the GP variants. **(C)** Representative western blot of CatL-treated and native VSV-GP(May 82V/544I), +/- PNGase F treatment, showing the glycosylation-dependent migration of GP (black arrows), GP_18K_ (open arrows), and GP_12K_ (white arrow).

Because GP1_18K_ appears to chase into GP1_12K_, the latter must be derived by cleavage of the former. Accordingly, we next sought to establish which GP1 sequences are lost upon GP1_18K_→GP1_12K_ cleavage. Previous work indicates that CatL-cleaved WT Mayinga GP1_18K_ comprises GP1 residues 33–190 (30). Making use of the N–linked glycan at position 40 GP1_18K_, the sole remaining glycan in this product (31), we analyzed the migration patterns of the VSV-GP(May 82V/544I) cleavage products prior to or following deglycosylation with Protein N–glycosidase F (PNGase F). For intact GP1ΔMuc and GP1_18K_ we observed a shift in the electrophoretic mobility of these products upon PNGase F treatment, whereas GP1_12K_ was unaffected (**Figure 7C**). Thus, N–terminal residues encompassing N40 are removed during GP1_18K_→GP_12K_ cleavage. Moreover, GP_12K_ was recognized by an antiserum raised against a peptide comprising GP1 residues 83–97 (13) **(Figure 7A)**, indicating that it contains this sequence. We propose, therefore, that the new N–terminus generated by GP1_18K_→GP_12K_ cleavage lies between GP1 residues 41 and 83. Whether GP1_18K_ also undergoes C–terminal cleavage remains unknown at present.

## Discussion

The 2013-2016 EBOV epidemic in West Africa led to an unprecedented number of infections and deaths but has enabled the first opportunity to examine how the virus evolves during prolonged human-to-human transmission. Viral sequences sampled during this epidemic revealed several non-synonymous mutations compared to the EBOV Mayinga reference strain, isolated in 1976 following limited human-to-human transmission (1–4). Two of these polymorphisms in GP—at positions 82 and 544—were previously shown to have opposing impacts on viral infectivity (15, 16). Here we show that these mutations affect distinct steps in EBOV entry, which points toward a novel mechanistic basis for the increased infectivity of the A82V mutation.

EBOV entry into cells depends on a number of host factors, including cysteine proteases CatB and CatL that act on GP to reveal the binding site (13, 14) for the universal filovirus receptor NPC1 (11, 12). Evidence suggests that these factors, while critical for entry, are insufficient to elicit GP membrane fusion (20, 32, 33) and that an additional host factor, acting as a fusion trigger, must exist.

Although viral fusion glycoproteins can be primed and triggered by many different factors, thermostability serves as a good general proxy for their capacity to undergo fusogenic conformational changes (23, 34, 35). We found that GPs bearing 544I were less thermostable than GPs bearing 544T, regardless of the residue at position 82, and that THL-cleaved GP exhibited incremental destabilization, with both 82V and 544I contributing to reduced stability versus their 82A and 544T counterparts. We propose that polymorphisms at position 544 primarily influence global GP stability, which may have ramifications for initial proteolytic cleavages in GP or even for fusogenic rearrangement. Due to its location in the GP2 internal fusion loop, residue 544 has been speculated to alter fusogenicity and/or bilayer insertion (10,

17, 18, 36). In GP peptides at low pH, 544I forms part of a “hydrophobic fist” together with 529L and 535F, which is proposed to be necessary for penetration of the host membrane (37). Although this study did not examine the effect of polymorphisms at position 544, we found no gross defect in fusion triggering or hemifusion in cells, only a significant delay in fusion kinetics between mutants bearing T or I at this position.

Although 544I has been demonstrated to increase infectivity in cell lines derived from different hosts, the promotion of infection by 82V occurs selectively (36), suggesting that its basis for enhanced infectivity is host species-dependent. Some evidence for 82V-induced instability lies in the mutant’s 3.47 resistance and decreased T_m_ values, but we argue that this is secondary to 82V’s role in facilitating GP cleavage by CatL. Cell type-dependent differences in CatL abundance or activity may explain the selectively heightened infectivity of 82V better than differences in NPC1 interaction.

The relative instability of GP bearing 82V has been cited as the primary source of its enhanced infectivity (10), but this mutation did not lead to increased probability of fusion triggering within cells, as might be expected with a less-stable protein. Instead, we saw that fusion triggering occurred more rapidly, as measured by overall lipid mixing rates and time-to-fusion following delivery to NPC1-positive compartments. We observed faster fusion kinetics for GPs bearing 82V, 544I, or a combination thereof. Although our live assays were unable to monitor the infection success of any given membrane fusion event, earlier triggering of viruses is generally associated with increased infectivity (38, 39). In addition to driving more rapid viral entry, the faster rates of fusion may enhance the probability of successful viral escape into the cytoplasm by facilitating evasion of the harsh conditions in the lysosome and/or lysosome-targeted host antiviral factors such as the late endosomal/lysosomal interferon-induced transmembrane proteins (IFITMs) (27, 40).

These observations are in contrast to *in vitro* proteolytic cleavage of GP by CatL, which is promoted specifically by the presence of 82V, but not 544I. GPs bearing 82V were both more labile, and intriguingly, they were processed to a novel GP1 cleavage product of approximately 12K in size (GP1_12K_). Our approximate mapping of this newly identified novel cleavage product indicates that the GP1_18K_ product usually obtained with WT GP undergoes further N–terminal cleavage by CatL to remove as many as 50 additional amino acid residues. This altered pattern of proteolytic cleavage was not observed following treatment with THL, but it may extend beyond CatL in cells, since a relative increase in infectivity was observed in U2OS cells lacking CatL.

Our findings suggest that residue 544 acts as a regulator of infectivity in cell culture by modulating overall GP stability and fusogenicity, whereas residue 82 more specifically influences proteolytic cleavage of GP by host endosomal cysteine cathepsins. This is reflected in increased relative infectivity under conditions at which cysteine cathepsin activity is depleted, either genetically or by chemical inhibition, as well as in increased proteolysis *in vitro* with the appearance of a novel cleavage product. Combining these effects affords an increase in the kinetics of fusion, explaining the previously observed increase in infectivity of the GP(82V/544I) double mutant.

We recently showed that a mutation near the N–terminus of GP1, R64A, inhibited viral membrane fusion and entry in a manner reversible by second-site mutations near the GP1 N–terminus (23). These second-site reversions were predicted to destabilize GP and enhance the exposure of GP1 N–terminal sequences to proteolysis. Our current findings argue that A82V accelerates viral membrane fusion and increases viral entry through a related mechanism—enhancing proteolytic cleavage at N–terminal sequences in GP1. Although more work is needed, both sets of observations are concordant with a hypothesis in which conformational changes at the GP1 base, coupled to NPC1 recognition by the receptor-binding site in the GP1 head, expose the base to further proteolytic cleavages that drives fusogenic rearrangement. We speculate that these GP1 N–terminal cleavages are normally mediated by the sequential activity of endosomal cysteine aminopeptidases such as CatC or CatH but can be more promiscuously carried out by cysteine endoproteases like CatL in the context of the A82V mutation, thereby contributing to the observed enhancements in viral entry and infection.

## Materials and methods

### Cells, viruses, and infection conditions

All infection experiments were carried out in Vero cells, cultured in Dulbecco’s modified Eagle medium (DMEM) (Life Technologies, Carlsbad, CA) supplemented with 2% fetal bovine serum (Atlanta Biologicals, Flowery Branch, GA), 1% penicillin-streptomycin (Life Technologies, Carlsbad, CA) and 1% GlutaMax (Life Technologies, Carlsbad, CA), or in U2OS cells cultured in McCoy’s 5A modified medium (Life Technologies, Carlsbad, CA), supplemented with 10% fetal bovine serum (Atlanta Biologicals, Flowery Branch, GA), 1% penicillin-streptomycin (Life Technologies, Carlsbad, CA) and 1% GlutaMax (Life Technologies, Carlsbad, CA). U2OS cells stably expressing monomeric NeonGreen fused to NPC1 (NPC1-mNG) were generated as previously described (26).

Pseudotyped VSVs bearing mutant EBOV GPs were generated as described previously (28, 41). Mutant Mayinga GPs were based on the EBOV/H.sapiens-tc/COD/1976/Yambuku-Mayinga isolate amino acid sequence (GenBank accession number AF086833). Mutant Makona GPs were based on the EBOV/H.sapiens-wt/LBR/2015/Makona-LIBR16393 (GenBank accession number KY744596). All GPs included a deletion of the mucin domain (Δ309–489) (42).

Prior to infection experiments, confluent Vero or U2OS cells were seeded in their respective growth media described above. VSVs bearing mutant GPs were diluted in the corresponding media before addition to cells. Infected cells were then maintained at 37°C for 14-16 h post infection before automatic counting of GFP-positive cells using a Cytation 5 Cell Imaging Multi-Mode Reader (BioTek Instruments, Winooski, VT) and using the onboard software for calculations of percentage infected cells.

### NPC1 domain C ELISA

Plates were coated with GP-specific mAb KZ52 (43) diluted to 2 µg/mL in PBS. VSV-GP mutants were normalized for GP content and then incubated at 37°C for 1 hour in the presence of 250 µg/mL thermolysin (Sigma-Aldrich, St. Louis, MO), before the addition of 10 mM phosphoramidon to stop the reaction. ELISA plates were blocked using PBS supplemented with 3% bovine serum albumin (PBSA; Thermo Fisher Scientific, Waltham, MA). THL-treated virus was added to blocked KZ52-coated plates and allowed to adsorb at 37°C for 1 hour. Following washing with 3% PBSA, a dilution series of FLAG-tagged NPC1-domain C (44) was added and allowed to bind for 1 hour at 37°C. Bound domain C was then detected using an HRP-conjugated anti-FLAG antibody (Sigma-Aldrich, St. Louis, MO) and Ultra-TMB substrate (Thermo Fisher Scientific, Waltham, MA). EC_50_ values were calculated using non-linear regression of data from two independent experiments with three technical replicates each.

### CA074 and 3.47 inhibitor experiments

Viral inhibitors 3.47 (Microbiotix, Worcester, MA) and CA-074 (Sigma-Aldrich, St. Louis, MO) were reconstituted in DMSO. Two-fold dilution series starting at 100 µM and 1 mM, respectively, were then made in complete DMEM (Life Technologies, Grand Island, NY) supplemented with 2% fetal bovine serum (Atlanta Biologicals, Flowery Branch, GA), 1% Penicillin-Streptomycin (Life Technologies, Carlsbad, CA) and 1% GlutaMax (Life Technologies, Carlsbad, CA). Diluted inhibitors were then added to confluent Vero cells, and uptake was allowed to proceed through 4 h incubation at 37°C. VSV-GP mutants were then diluted 1:1,000 in complete DMEM and added to inhibitor-treated cells. At 14-16 h post-infection, cells were fixed in 4% PFA and stained with Hoechst nuclear stain before GFP-positive cells were counted using a Cytation 5 Cell Imaging Multi-Mode Reader (BioTek Instruments, Winooski, VT) with the onboard software for calculations of percentage infected cells.

### In vitro proteolysis

VSV-GP mutants, normalized for GP content, were incubated with activated human recombinant CatL (R&D systems, Minneapolis, MN) at 2 ng/µl or THL (Sigma-Aldrich, St. Louis, MO) at 250 µg/mL. Cleavage by CatL was allowed to proceed at 37°C for 15, 30, 60, 90, 120, 150 or 180 min before the reaction was stopped by addition of 0.1 mM E-64. Cleavage by THL was allowed to proceed at 37°C for 30, 60, or 120 min before the reaction was stopped by addition of 10 mM phosphoramidon. Cleavage products were then deglycosylated through treatment with 250 U PNGase F (New England Biolabs, Ipswich, MA) for 16 h at 37°C. Parallel samples of VSV-GP(May 82V/544I) treated with CatL for 60 min were also left without deglycosylation for the analysis of the presence of the glycan at N40.

### SDS-PAGE and western blot

Deglycosylated and native samples containing GP cleavage products were analyzed by SDS-PAGE using 10% Tricine protein gels (Thermo Fisher Scientific, Waltham, MA). Samples were transferred onto 0.2 μm pore size Protran nitrocellulose membranes (Sigma-Aldrich, St. Louis, MO) followed by western blot for GP1 using a rabbit polyclonal sera recognizing a peptide (TKRWGFRSGVPPKVV) overlapping the receptor-binding site. IRDye 680LT Goat anti-Rabbit IgG 680 secondary Ab (LI-COR, Lincoln, NE) was used at a dilution of 1:10,000, and the final blot was then imaged using a LI-COR Fc fluorescent imager.

### GP thermostability assay

GP thermostability assay was conducted as previously described (23). VSVs bearing mutant GPs were treated with THL at 500 µg/mL for 30 min at 37°C, or mock treated, before the addition of 10 mM phosphoramidon to stop the reaction. Cleaved and uncleaved virus were diluted in PBS and then heated at a range of temperatures spanning 42 to 64°C (cleaved) and 46 to 68°C (uncleaved) for 10 min followed by cooling to 4°C using a thermal cycler (Applied Biosystems, Foster City, CA). After cooling, virus was directly captured onto high-binding 96-well half-area ELISA plates (Corning, Corning, NY). Plates were then blocked using 3% bovine serum albumin in PBS. GP was detected using KZ52, a conformation-specific anti-EBOV GP monoclonal antibody. Antibody bound to GP was then detected with anti-human antibody conjugated to HRP (EMD Millipore, Burlington, MA) and Ultra-TMB substrate (ThermoFisher, Grand Island, NY). All binding steps were carried out at 37°C for 1 h. Binding curves were generated using Prism (non-linear regression, variable slope [four parameters]; GraphPad Software, La Jolla, CA). T_m_ were calculated from two independent experiments, each with three replicates.

### Virus labeling for live-cell microscopy

Purified virus (1 mg/ml) was labeled with 50 µM 1,1’-dioctadecyl-3,3,3’,3’-tetramethylindodicarbocyanine (DiD) dye while being agitated for 1 h at 4°C. Excess dye was removed by ultracentrifugation of the virus through a 10% sucrose cushion for 2 h at 107,000 × *g* and 4°C using an SW41 rotor (Beckman Coulter). Labeled virus pellets were resuspended at a viral protein concentration of 1 mg/ml, aliquoted, and stored at -80°C until use.

### Live imaging

Live-cell microscopy was performed as previously described (27) with an AxioObserver.Z1 widefield epifluorescence microscope (Zeiss, Oberkochen, Germany) equipped with a 40×/ 1.3 N.A. objective, DAPI/GFP/Texas Red/Cy5 filter set and heated environmental enclosure maintained at 37°C. U2OS cell monolayers were seeded onto fibronectin-coated 35-mm glass coverslip dishes (MatTek, Ashland, MA) 24 h before experiments. Cells were chilled for several minutes on ice before spinoculation of DiD-labeled virus onto monolayers at 1,500 × *g* and 6°C for 20 min. Unbound particles were removed by five washes with cold PBS, and 500 μl cold imaging buffer (140 mM NaCl, 2.5 mM KCl, 1.8 mM MgCl_2_, 20 mM HEPES, 5 mM sucrose, 2 μM Hoechst 33342, and 2% FBS) was added to cover the cells. The dish was immediately mounted on the microscope objective and focused. The coverslip dish was then flooded with 1.5 ml warm imaging buffer to mark the start of experiments (*t* = 0). Images were acquired every 10 s over the duration of the experiments using a single Z-section, which encompassed nearly all cell-associated particles.

## Data analysis

Image analysis and single-particle tracking were performed using Volocity software (PerkinElmer, Waltham, MA) as previously described (26). Image files were not manipulated, apart from minor adjustments in brightness and contrast. Viral puncta were thresholded by initial intensity and size. Puncta falling outside the range of 0.25 to 1 μm^2^ expected of individual DiD-labeled virions were excluded from single-particle analysis. Virions were considered colocalized with NPC1 or Rab5 only if the cellular marker punctum exceeded the background signal by 30% or more and if the intracellular and viral puncta co-trafficked with greater than 70% overlap of signals. Mean measurements (± s.d.) were derived from three separate experiments, unless otherwise indicated.

## Acknowledgements

We thank I. Gutierrez, E. Valencia, L. Polanco, C. Harold, and T. Krause for laboratory management and technical support. We also thank M. Ng for providing reagents and for exploratory work related to this project. We acknowledge funding support from NIH R01 AI134824 (to K.C.).

